# VARIATION OF MULTIPLE PLANT DEFENSES AND HERBIVORY REVEAL DIFFERENT PATTERNS OF ADAPTATION ACROSS A PRODUCTIVITY GRADIENT

**DOI:** 10.1101/2025.10.31.684923

**Authors:** Jacob E. Herschberger, Eduardo S. Calixto, Valerie R. Campbell, John L. Maron, Philip G. Hahn

## Abstract

Plants with large geographic distributions experience varying biotic and abiotic conditions across their range that can influence plant defense traits among populations. Whether populations inhabiting warmer, wetter and more productive regions face greater pressure and results in higher investment in defense compared to populations from cooler, drier, and less productive regions remains uncertain. Here we quantified defense traits and herbivory on *Solanum carolinense L.* across a large climate and productivity gradient. We sampled populations spanning the entire north-south range (29–44° N) of *S. carolinense*, (33 populations; ∼15 plants per population) and employed a common garden experiment (15 populations; ∼5 plants per population). We examined whether climate predicts 1) plant defense traits (specific leaf area, trichome density and glycoalkaloid concentration), 2) herbivore damage, and 3) the correlation of traits and herbivore damage in field and common garden plants. Trichome density and herbivore damage were higher for *S. carolinense* at the center of its range, while glycoalkaloid concentrations were negatively associated with warmer, wetter climates both in the field and in the common garden. In the field, plants with higher glycoalkaloid production experienced reduced herbivory, while in the common garden plants with greater SLA received more damage. Overall, these findings suggest that different types of defensive traits within a single species may follow different ecological and evolutionary trends and highlight the need for trait-specific considerations when applying plant defense hypotheses at the intraspecific level.

## Introduction

Many plant species have large geographic distributions where different populations experience divergent abiotic and biotic conditions (Abdala-Roberts et al., 2016; López-Goldar and Agrawal, 2021). Adaptation to particular abiotic and biotic conditions can result in the evolution of unique physiological or defense traits among widely dispersed populations (Bickford, 2016; Agrawal et al., 2022). These phenotypic changes across geographical ranges can also be due to plasticity, or both plasticity and genetic differentiation (López-Goldar and Agrawal, 2021; Agrawal et al., 2022). A substantial body of work has shown these mechanisms at work, creating population differentiation in both plant physiological and life history traits (Endara and Coley, 2011; Massad et al., 2011). However, environmental gradients do not always produce population differentiation in defense traits, and whether this is because herbivore pressure does not vary predictably across these gradients, or for other reasons is not entirely clear (Maron et al., 2014; Louthan et al., 2015; Anstett et al., 2016).

Herbivory is typically expected to increase across climate and productivity gradients, because warmer, wetter climates results in greater plant productivity and therefore greater herbivore densities (Woods et al., 2012; Hahn et al., 2019; Agrawal et al., 2022; Croy et al., 2022). Empirical studies and meta-analyses have demonstrated that herbivory is often positively associated with gradients in latitude (Pennings and Silliman, 2005; Pennings et al., 2009; Salazar and Marquis, 2012; Zvereva and Kozlov, 2021), climate (independent of latitude) (Hahn et al., 2019, 2021) or elevation (Galmán et al., 2018, 2019; Kozlov et al., 2022; Zvereva et al., 2022), yet these general trends are not always supported (Anstett et al., 2014; Moreira et al., 2015, 2018; Kooyers et al., 2017; Agrawal et al., 2022). Some studies have reported non-linear trends in herbivory across a productivity gradient that can be influenced by the density of the host plant species which can be more abundant at the range center (Woods et al., 2012; Agrawal et al., 2022) or the more favorable edge (Fagan and Bishop, 2000). In general, herbivore abundance and how it is affected by overall community productivity and host plant abundance likely determines underlying spatial patterns of herbivory (Crutsinger et al., 2013; Anstett et al., 2014, 2016; Baskett and Schemske, 2018).

Although one might expect plant defenses to vary in concert with herbivore pressure across a species’ distribution, both theoretical predictions and empirical results are mixed (Moles et al., 2011; Anstett et al., 2016; Hahn and Maron, 2016). A set of general hypotheses assert that increasing herbivore pressure, for example towards the equator (Pennings et al., 2009), in more productive areas (Hahn and Maron, 2016; Kooyers et al., 2017; Hahn et al., 2019), or in the center of a geographic range (Anstett et al., 2016; Agrawal et al., 2022), should lead to greater plant defense in those areas. An opposing prediction is that intraspecific plant defenses should be greater in low-resource/high-stress environments, which has also been supported by numerous studies (Robinson and Strauss, 2018; Croy et al., 2022; Calixto et al., 2025). Plant growth is greater in more productive regions; therefore, plants may be able to outgrow herbivory or replace plant tissue at higher rates compared to plants in slow growth environments. This suggests that, in more abiotically stressful environments, defenses which reduce herbivory might be more valuable compared to more productive environments (Coley et al., 1985; Hahn and Maron, 2016). Thus, plant defense trends across a species’ range might differ depending on whether they are driven by herbivory levels or the resource availability for plant growth potential.

In this study, we quantify multiple defense traits (specific leaf area, trichome density and glycoalkaloid concentration) and herbivore damage for *Solanum carolinense* L. across a climatic gradient. We used the entire south-north (29-44^°^ N) species range to disentangle plant defense-herbivore associations. We coupled the field observations with a common garden experiment at the southern range edge (29^°^ N) to separate genetic and environmental effects on traits and herbivory. Using a piecewise structural equation model (pSEM), we took a holistic approach that addressed the following questions: 1) Do herbivore damage and plant defenses have a linear or non-linear association with climate? 2) Which defense traits are correlated with herbivore damage. And 3) Do herbivore damage and defense trait trends persist when plants are grown in a common garden? This study enabled us to better understand plant-herbivore interactions across spatial distributed populations.

## Materials and Methods

### Study species and populations

*Solanum carolinense* L. is an herbaceous perennial plant native to the eastern United States (29-44^°^ N and 70-98^°^ W; Fig. 1A). It grows in disturbed areas such as lawns, roadsides, and crop fields (Solomon, 1986; Kariyat et al., 2013). Flowering occurs mostly in June and July, and fruiting occurs in August and September (Bassett and Munro, 1986; Kariyat et al., 2013). *Solanum carolinense* is a close relative of crop species such as *S. lycopersicum* L. (Tomato), *S. melongena* L. (Eggplant), and *S. tuberosum* L. (Potato). Plants in the genus *Solanum* produce multiple different glycoalkaloids that function as a defense against various herbivores (Chowański et al., 2016; Zhao et al., 2021; Hauri et al., 2024). The two dominant glycoalkaloid compounds produced by *S. carolinense* are α-solasonine and α-solamargine, which are effective in reducing leaf feeding, and negatively impacting insect herbivore performance and fecundity (Wierenga and Hollingworth, 1992; Cipollini and Levey, 1997a, b; Güntner et al., 2000; Chowański et al., 2016). *Solanum carolinense* also has physical herbivore defenses including prickles and trichomes that are effective in protecting the plant against larger insect and non-insect herbivores (Kariyat et al., 2013, 2017; Nihranz et al., 2019). Lastly, this plant can tolerate some levels of herbivory by compensatory regrowth and asexual rhizomatous growth (Wise and Mudrak, 2021). Common insect species that feed on this plant across its range are leaf beetle herbivores (Coleoptera: Chrysomelidae) such as *Leptinotarsa juncta* G. (False Colorado Potato Beetle), *Epitrix fuscula* C. (Eggplant Flea Beetle), and *Gratiana pallidula* B. (Eggplant Tortoise Beetle) (Wise, 2007 & personal observations). Leaf herbivory is also less frequently inflicted by herbivores such as *Tildenia inconspicuella* M. (Eggplant Leaf Miner), *Gargaphia solani* H. (Eggplant Lace Bug), *Prodiplosis longifila G.* (Citrus Gall Midge), and sometimes the generalist herbivore *Manduca sexta* L. (Tobacco Hornworm) (Wise, 2007), *Spodoptera ornithogalli* (Guenée) as well as other generalist herbivores.

**Figure 1.**
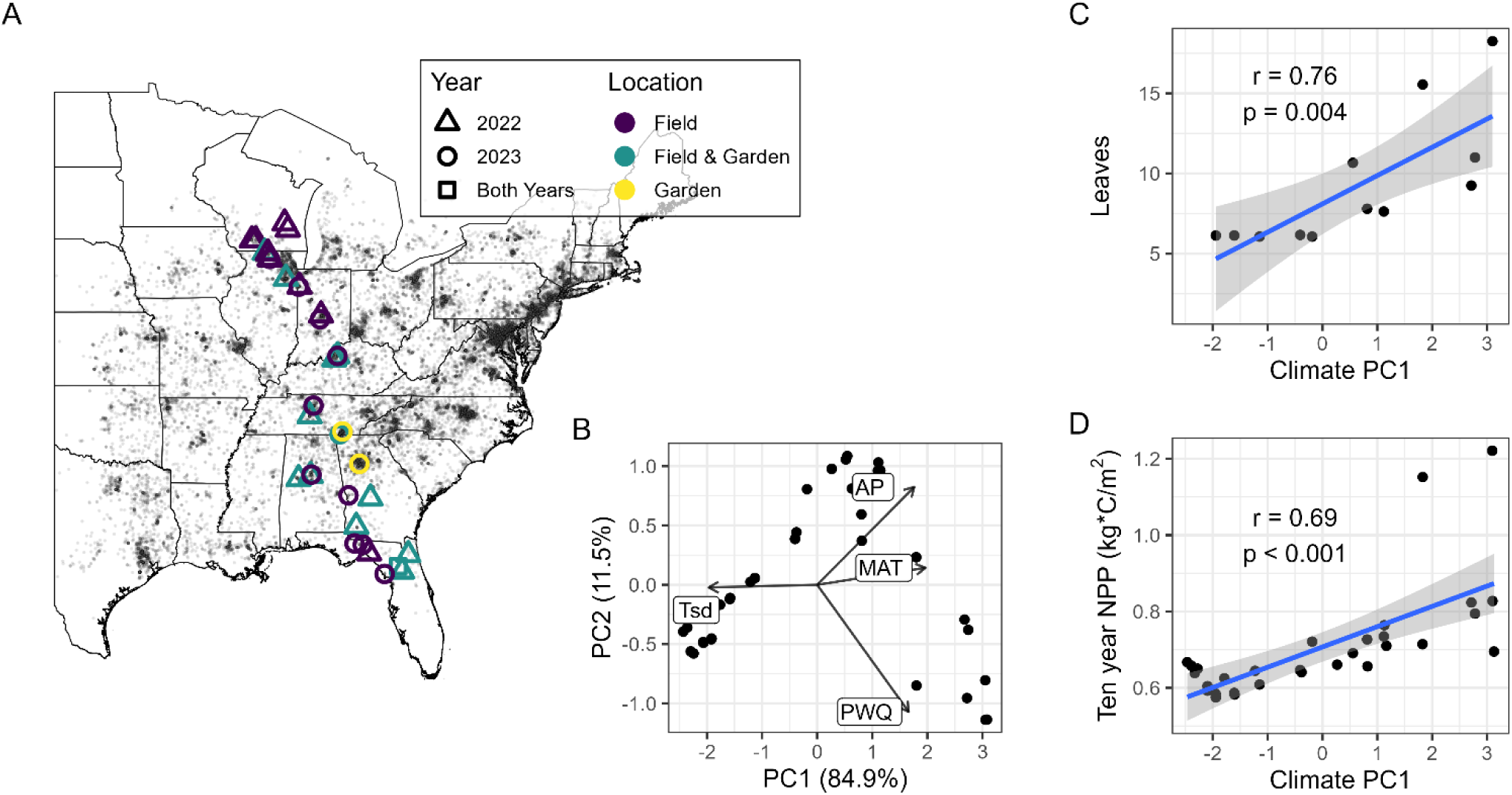
A) A map of *Solanum carolinense* plant populations surveyed in the field or collected for the common garden (large points) and iNaturalist observations (small points). Also shown is B) biplot of the principal component analysis and the first axis of the climate PCA plotted against C) plant size, measured as number of leaves, at a subset of populations from the field surveys in 2022, and D) 10-year average annual NPP from MODIS satellite images.

We selected a south-north transect across the entire *S. carolinense* native range spanning approximately 15^°^ in latitude, from 29-44^°^ N (Fig. 1A). Mean annual temperatures and precipitation across this range vary from 8-21^°^C and from 908-1508 mm, respectively. During summers of 2022 and 2023 we sampled 32 populations across the geographic gradient. We sampled *S. carolinense* populations that contained at least six ramets more than two meters apart. Twenty-one populations were sampled in 2022, 11 populations were sampled in 2023, and one population was sampled in 2022 and 2023 (n = 33 surveys total).

### Climate and NPP extraction

The rgee package (version 1.1.8) in R (version 4.5.3) was used to interact with the Google Engine image collections API and extract 50 year average, 1 km resolution (WORLDCLIM/V1/BIO) and 8-day Net Primary Productivity (NPP) (MODIS/061/MOD17A2HGF) at the *S. carolinense* field locations (Hijmans et al., 2005; Running & Zhao 2019). We specifically extracted bioclim variables such as the mean annual temperature (MAT), annual precipitation (AP), standard deviation of yearly temperature (Tsd), and the precipitation of the warmest quarter (PWQ). We chose MAT, AP, and PWQ since temperature and precipitation can both increase primary productivity across terrestrial ecosystems (Del Grosso et al., 2008). Additionally, Tsd was found to be influential on the production of chemical defenses across a latitudinal gradient (Anstett et al., 2018). We also extracted the annual NPP for a 10-year period (2013-2022) within a 5km radius of the plant populations and averaged across the 10 years. We selected the 5km radius buffer because some of the plant populations were in suburban areas and satellite sensors were not able to get the appropriate NPP within smaller buffers around the plant population.

To reduce the dimensionality of the four selected climate variables, we scaled (mean = 0, SD = 1) the variables and performed a principal component analysis (PCA) using the princomp function in the base R stats package (R Core Team, 2022). The first climate PC axis explained 84.9% and the second axis explained 11.5% of the variation (Fig 1B), where productivity-associated climate variables (AP, MAT, and PWQ) loaded positively on the first axis, and Tsd loaded negatively on first axis (Fig. 1B). Lastly, climate PC1 had a strong correlation with satellite derived 10-year average annual NPP (r = 0.69, P < 0.001, Fig. 1D). We used the first climate PC axis because it captures aspects of climate variation relevant to productivity and plant-herbivore interactions.

### Field population sampling

#### Herbivore damage

To determine herbivore damage across the geographic gradient, we sampled 5-30 individuals within each source population during June through early August in 2022 and 2023 (n= 521 and mean plants per population = 15.8 ± 9.7 SD). We haphazardly selected mature (i.e., budding or blooming) plants further than two meters apart. We chose mature plants, so the observed herbivore damage is a good representation of cumulative herbivory that occurred throughout the growing season and is also when herbaceous plants switch from defending leaves to allocating resources to reproduction (Boege and Marquis, 2005; Rusman et al., 2020). Herbivore damage was quantified by inspecting each leaf on the plant and visually estimating the total average percent damage of the whole plant (Robinson et al., 2023). Since *S. carolinense* is relatively small (20-40 cm tall) and typically only produces 10-50 leaves per plant, we could easily visually estimate herbivory of the whole plant. We also measured plant size as the number of leaves during the 2023 surveys, which was strongly correlated with the Climate PC1 axis (Figure 1C). During the surveys, we regularly observed the specialist herbivores (i.e., *Leptinotarsa juncta*, *Epitrix fuscula*, and *Gratiana pallidula*, personal observations) and other generalist herbivores (e.g., *Spodoptera ornithogalli*, aphids, grasshoppers, white flies, etc.). Although we tended to notice these more frequently in the center of the range (Table S1), these observations were too sparse to formally analyze.

#### Plant defense traits

To determine how plant defense traits varied across latitudes, we measured trichomes and glycoalkaloids on the same plants where herbivory was quantified. We also measured specific leaf area (SLA), which provides a leaf growth construction metric and can indicate plant palatability status (Schädler et al., 2003; Agrawal and Fishbein, 2006). We collected the two top, fully developed leaves from each of the plants and oven-dried them at 50^°^C for 48 hours. Trichome density was estimated by counting the number of trichomes in a 0.24 cm^2^ circle randomly placed between secondary leaf veins. We used a digital USB microscope (Aven 26700-209-PLR) at 5X magnification to photograph and count trichomes on the leaves. Specific leaf area (SLA; leaf area/dry mass) was measured by cutting a leaf piece that avoided the midrib and major secondary leaf veins, measuring the area to the nearest 0.1 mm^2^, and then weighing the leaf to the nearest 0.1 mg. We measured the leaf area by first scanning the cut leaves with a Cannon scanner (K10486) and then used ImageJ to measure the leaf area.

Lastly, total glycoalkaloids were measured using a direct hydrolysis-colorimetric protocol, modified slightly from previous studies (Cipollini and Levey, 1997a; Walls et al., 2005). Although, this method does not quantify specific glycoalkaloids, *S*. *carolinense* only produces two glycoalkaloid variants (α-solasonine and α-solamargine); both are effective in defending against herbivores (Wierenga and Hollingworth, 1992; Güntner et al., 2000; Chowański et al., 2016), and this protocol has been commonly used in previous studies that quantify total glycoalkaloids in *S. carolinense* (Cipollini and Levey, 1997b, a; Walls et al., 2005). We scaled the glycoalkaloid quantification protocol to use a microplate reader rather than a cuvette reader that had been used in previous protocols. First, we weighed and ground 20 ± 2 mg of leaf material to a fine powder with three 1.4 mm Fisherbrand ceramic beads in 2 ml microcentrifuge tubes. Then we added 0.5 ml of 0.5 M HCl and incubated the sample in a 100°C water bath for two hours. Next, we neutralized the solution with 0.5 ml of 0.5 M NaOH and added 0.5 ml acetic acid. After centrifuging the solution for five minutes at 12000 g, we transferred 0.9 ml of the supernatant to a new 2 ml tube and added 0.6 ml DI H_2_O. We then used 0.5 ml of the previous sample and added 0.5 ml acetate buffer (5.44 g sodium acetate, 100 ml DI H_2_O, and 2.4 ml acetic acid; pH 4.7), 0.1 ml methyl orange, and 0.5 ml methylene chloride. After mixing this solution for three minutes and allowing to settle for three minutes, we pipetted 200 ul of the infranatant into a u-shaped, glass-bottomed 96 well microplate and measured the absorbance at 420 nm using a microplate reader (Multiskan^TM^ SkyHigh, Thermo-Scientific, United States).

### Common garden study design

To examine genetic variation in defensive traits and how these traits translate to variation in herbivore damage, we grew *S. carolinense* in a common garden near the southern range edge in an old-field site on the University of Florida campus (Gainesville, Florida USA; lat. 29.626799, long. -82.356232) from April to August 2023. We used roots collected from ∼5 plants 2 m apart within each population during 2022 (n= 70 plants total from 15 populations, and mean plants per population = 4.7 ± 1.5 SD). We cut the roots to 0.5-1 g each before planting the cuttings in 3.8 L pots that contained a 1:1 mixture of topsoil and potting mix. Plants were started in the greenhouse on April 19, 2023, in a temperature range of 21-32° C, a photoperiod of approximately 13 L:11 D, and 60% relative humidity. After 4 weeks of growth, the plants were transferred to the outdoor common garden and placed 30 cm apart in a randomized block design on wooden 1.2 m x1.2 m pallets. Surrounding vegetation was occasionally mown and plants were occasionally watered. We then observed herbivore damage every 2-3 weeks during the growing season. The most common herbivore in the common garden was larvae of the generalist *Spodoptera ornithogalli* and we did not observe any of the specialist beetles of *S. carolinense*. We used the average herbivory data from early June to late July. This timeline provides good representation of the herbivory across the growing season (Fig. S1). Additionally, we quantified SLA, trichomes, and glycoalkaloids in the same manner as in the field surveys. The leaves to measure the defense traits were collected at the end of the growing season on August 17, 2023, to reduce the effects of harvesting leaves on herbivory and trait expression.

### Data and statistical analysis

We used a piecewise structural equation model (pSEM) to test herbivory-defense relationships across the productivity gradient, which allowed us to simultaneously evaluate all relevant relationships between herbivore damage, plant traits, and how they vary across the productivity gradient. We used the psem function from the piecewiseSEM R package, which provides the option to include random effects and various generalized linear models (Lefcheck, 2016). For our main response (endogenous) variable, proportion of herbivore damage, we used the ordered beta distribution, which is appropriate for continuous proportions including 0 and 1 (Kubinec 2023). We log-transformed trichomes, SLA, and Glycoalkaloids, to improve the normality of the residuals. Additionally, after transformations all the variables, except herbivory, were centered and scaled (mean = 0, SD = 1) to increase model stability (Schielzeth, 2010).

The structure of our pSEM to test our first two hypotheses regarding how climate and traits predict 1) herbivory and 2) how climate predicts traits, was as follows: the first sub-model included herbivore damage as the response and climate PC1 as the predictor. Plant traits were also included as predictors to infer the relationship between herbivory and plant defense traits. To test whether plant defense traits are related to the climate PC1, additional sub-models were included where each defense trait was a response variable and climate PC1 as the predictor to test for clinal variation in each trait. In the field, the random effects were modeled where populations were nested within year. The pSEM for the common garden was identical except population was the only random effect to account for the multiple plants within each population.

To explicitly test the linear vs non-linear trends of herbivory and plant traits in our first hypothesis, we build two versions of the pSEM models described above: one version included climate PC1 as only a linear term and the second also included the quadratic term for the climate PC1 variable (in all sub-models). This was done for both the field and common garden data.

We used the psem and summary functions from the piecewiseSEM package, respectively, to build the model and extract the statistics (i.e. coefficients, p-values, and fit parameters) of the model (Lefcheck, 2016). Because all predictors were centered and scaled prior to analysis, the raw model estimates are interpretable as standardized path coefficients (Lefchek 2016). For the herbivory sub-model, which used an ordered beta distribution (Kubinec 2023), coefficients represent the change in herbivory on the logit scale per one standard deviation change in each predictor. We assessed overall model fit using Fisher’s C and p-value. We also calculated the variance explained by the fixed effects (R^2^m), and the variance explained by the full model (R^2^c) using the r() function in performance package (Lüdecke et al., 2021). For indirect effects (e.g., the effect of climate PC1 on herbivory mediated through defensive traits), we multiplied the coefficients involved in the mediated path (Grace et al. 2012).

Lastly, to test our third hypothesis regarding the correlation of plant traits between the field and garden plants, we used the cor.test function in the R base package to test the correlation of the field and garden mean population trichome, glycoalkaloid, and SLA values.

## Results

### Results from field population surveys

To explore how climate potentially influenced field-measured plant defense traits across the range of *S. carolinense* and how these traits link to herbivore damage we used a pSEM model with quadratic terms for climate (AIC = 2614) because it was a better fit than a linear model (AIC = 2623). This model fit the field data well (Fisher’s C = 3.7, df = 4, p = 0.45), after correcting for directed separation between SLA and trichomes (r = -0.24, p < 0.001, Table 1). There was a hump-shaped quadratic relationship between the climate PC and trichomes, where trichome density was higher at the center of the plant range (i.e., low to mid values of the climate PC; Figs. 2A & 3A). Glycoalkaloid concentrations decreased linearly with the climate PC (Figs. 2A & 3B). There was no linear or quadratic trend between SLA and the climate PC (Table 1). Additionally, herbivore damage had a hump-shaped relationship with the climate PC (Figs. 2A & 3C) and was negatively correlated with glycoalkaloids (Figs. 2A & 3D) but did not correlate with SLA or trichomes (Fig. 2A & Table 1). The indirect effects of the climate PC on herbivore damage, mediated through glycoalkaloids or trichomes, were weak relative to the direct effects (Table S2).

**Figure 2.**
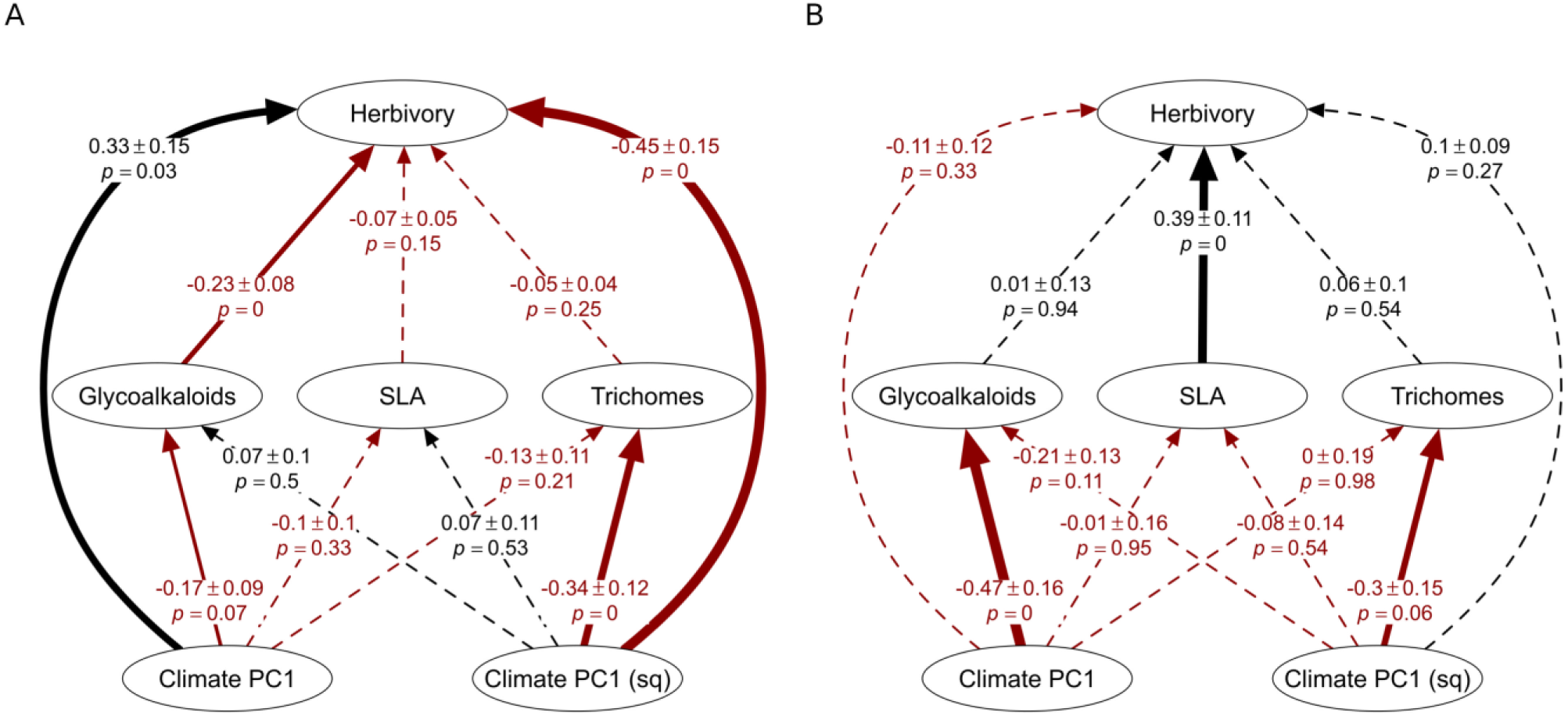
The pSEM diagrams of A) field and B) garden results. Solid lines relationship between the predictor and the response (*P* < 0.1), and the dashed lines indicate a non-significant relationship. Black lines indicate a positive relationship, and red lines indicate a negative relationship. The widths of the lines indicate the strength of the at least marginally significant relationships. Coefficient for the standardized variables, plus/minus the SE and p values are shown for the paths.

**Figure 3.**
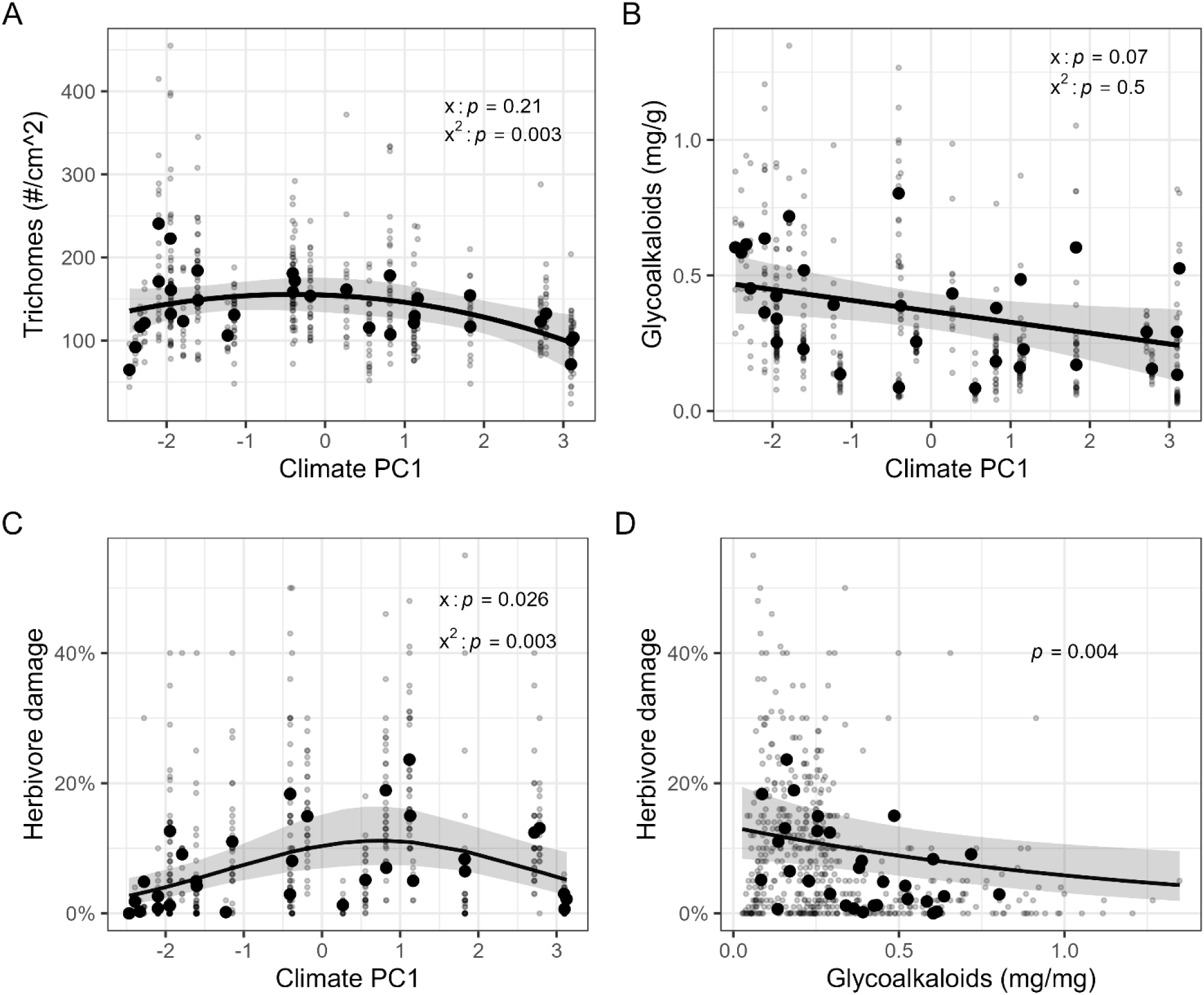
Graphs from field surveys of A) trichomes, B) glycoalkaloids and C) herbivory in relation to the climate PC1 axis. D) shows herbivory in relation to glycoalkaloids. Small black data points indicate individual plants, and large points indicate population averages. The lines indicate significant generalized quadratic/linear trend of the relationship between the predictors and responses of individual data points, with the shading representing the 95% confidence intervals. Herbivory is fitted with an ordered beta distribution.

**Table 1.** Full model coefficients for field and garden pSEM. Paths with *P* < 0.1 are bolded.

|  |  | Field |  |  |  |  | Common Garden |  |  |  |  |
| --- | --- | --- | --- | --- | --- | --- | --- | --- | --- | --- | --- |
| Response | Predictor | Estimate | SE | P | R<sup>2</sup>m | R<sup>2</sup>c | Estimate | SE | P | R<sup>2</sup>m | R<sup>2</sup>c |
| **herbivory** | **ClimPC1** | **0.332** | **0.149** | **0.026** | 0.20 | 0.58 | -0.112 | 0.116 | 0.334 | 0.27 | - |
| **herbivory** | **ClimPC1<sup>2</sup>** | **-0.454** | **0.151** | **0.003** |  |  | 0.096 | 0.088 | 0.274 |  |  |
| herbivory | Trichomes | -0.073 | 0.051 | 0.146 |  |  | 0.063 | 0.103 | 0.539 |  |  |
| herbivory | SLA | -0.073 | 0.050 | 0.146 |  |  | **0.388** | **0.105** | **0.000** |  |  |
| **herbivory** | **Glycoalkaloids** | **-0.232** | **0.080** | **0.004** |  |  | 0.011 | 0.13 | 0.935 |  |  |
| Trichomes | ClimPC1 | -0.132 | 0.105 | 0.210 | 0.14 | 0.42 | -0.005 | 0.186 | 0.979 | 0.13 | 0.48 |
| **Trichomes** | **ClimPC1<sup>2</sup>** | **-0.340** | **0.115** | **0.003** |  |  | **-0.296** | **0.155** | **0.055** |  |  |
| SLA_s | ClimPC1 | -0.101 | 0.104 | 0.332 | 0.01 | 0.58 | -0.011 | 0.164 | 0.945 | 0.01 | 0.22 |
| SLA_s | ClimPC1<sup>2</sup> | 0.070 | 0.109 | 0.527 |  |  | -0.084 | 0.136 | 0.537 |  |  |
| **Glycoalk** | **ClimPC1** | **-0.166** | **0.092** | **0.070** | 0.03 | 0.70 | **-0.471** | **0.158** | **0.003** | 0.26 | 0.52 |
| **Glycoalk** | ClimPC1<sup>2</sup> | 0.065 | 0.097 | 0.502 |  |  | -0.213 | 0.132 | 0.107 |  |  |
| **SLA ~** | **~Trichomes** | **-0.240** | - | **0.000** | - | - | **-0.244** | - | **0.022** | - | - |

### Results from common garden study

To explore how climate influences genetically based differences in plant defense traits and how they link to herbivore damage, we used a pSEM model with quadratic terms for climate (AIC = 409.3) for consistency with the field results, although the linear pSEM fit similarly (AIC = 408.4). The quadratic psem model fit the common garden data well after accounting for directed separation between trichomes and SLA (Fisher’s C = 4.06, df = 4, p = 0.40). Overall, we found that there were similar trends between plant defense traits and the climate PC, compared to the field. Trichomes had a hump-shaped quadratic relationship (Figs. 2B & 4A) and glycoalkaloids had a negative relationship with the climate PC (Figs. 2B & 4B), whereas there was no relationship between SLA and the climate PC (Table 1). In contrast to patterns in the field, there was no relationship between herbivore damage and the climate PC in the common garden (Fig. 2B). Lastly, there was no relationship between glycoalkaloids or trichomes and herbivore damage, although there was a positive correlation between damage and SLA (Fig. 4D).

**Figure 4.**
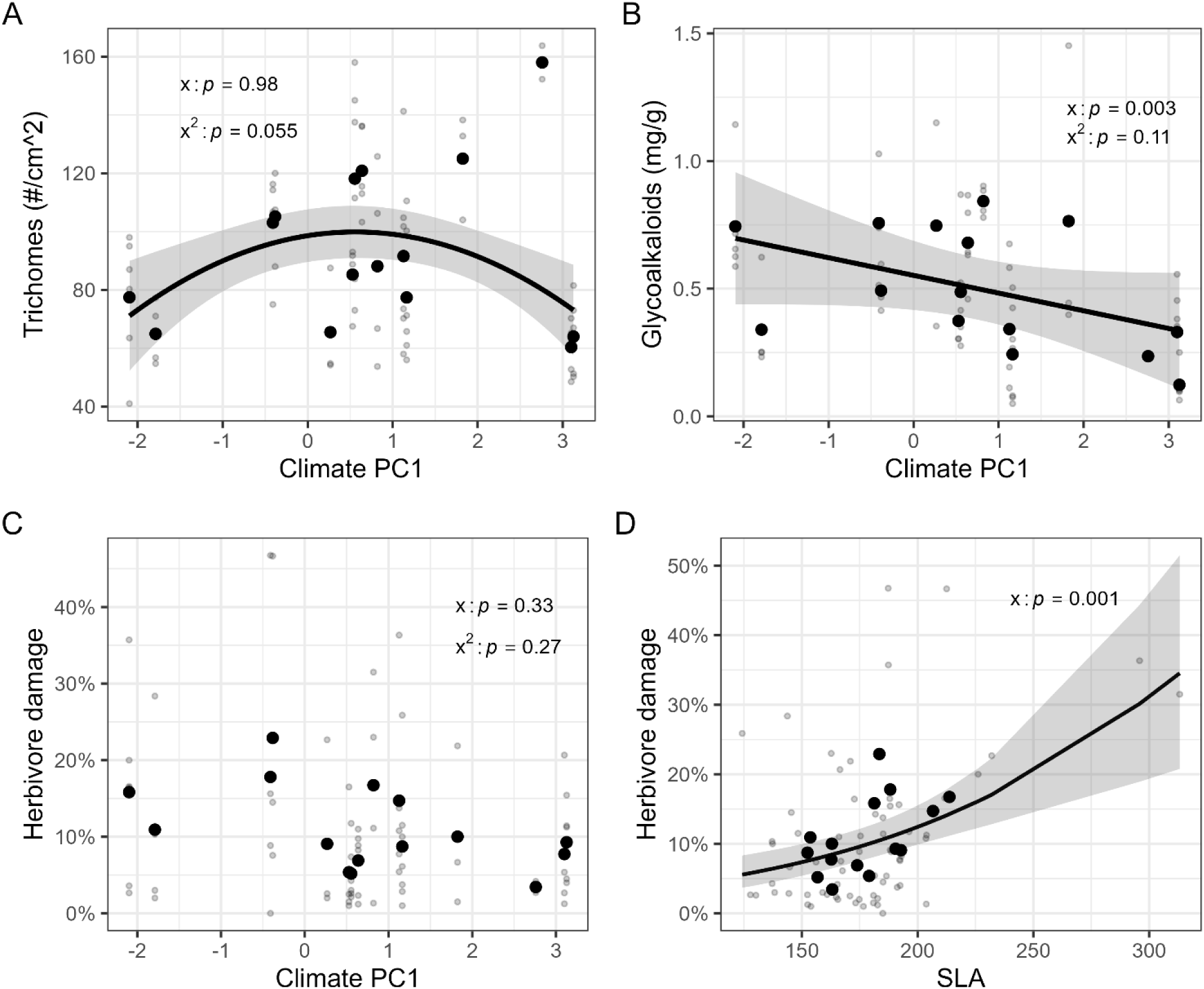
Garden data plots of A) trichomes, B) glycoalkaloids, and C) herbivore damage in relation to the climate PC. D) shows the relationship between herbivore damage and SLA. Transparent grey data points indicate individual plants, and solid black data points indicate population averages. The lines indicate significant generalized quadratic/linear trend of the relationship between the predictors and responses of individual data points, with the shading representing the 95% confidence intervals. The two points with extreme SLA values in D do not affect the estimated relationship or significance.

## Discussion

Intraspecific studies testing whether there is an association between climate and productivity gradients and plant defense/herbivore damage have had mixed conclusions (Moles et al., 2011; Woods et al., 2012; Moreira et al., 2018). However, previous studies have typically only measured single defensive trait and across only a portion of a species’ full distribution. To bolster existing knowledge, we quantified the association between plant productivity, herbivore damage, and multiple defense traits across populations that span the full south-north distribution of *Solanum carolinese* in both the field and a common garden. We found that defensive traits follow different patterns across a productivity gradient. Glycoalkaloid concentration had a negative association with the productivity gradient and trichomes density was the highest at the center of the gradient in both the field and common garden. Moreover, while higher glycoalkaloid concentrations correlated with reduced herbivory in the field, this trend did not persist in the common garden, maybe due to different herbivory types and intensity. Importantly, the results reveal different patterns between defense traits but consistent patterns in the field and common garden, demonstrating that some plant defenses may be influenced by herbivore pressure while other defense traits may be more influenced by climate and productivity.

Trichomes were produced at higher densities in the middle of the climate gradient, which corresponds to the center of the *S. carolinense* range, with significantly lower densities toward both northern and southern range edges (Figs. 2, 3A, & 4A). These results support the range center hypothesis where plant defense production coincides with higher specialist herbivore abundance at the center of a plant species range (Anstett et al., 2016). Specialist herbivores are typically most abundant and diverse at the center of the host plant range due to better connectivity among host plants and larger plant population sizes (Crutsinger et al., 2013; Anstett et al., 2016). The main herbivores of *S. carolinense* are specialist chrysomelid beetle species, *Leptinotarsa juncta*, *Epitrix fuscula*, and *Gratiana pallidula*, which are more commonly found at center latitude populations and might impose selective pressure on *S. carolinense* trichomes (personal observation; Table S1). The weak but consistent association between trichome density in the field and common garden experiment suggests that trichome patterns found across the range may have a genetic basis (correlation of population means: r = 0.40, df = 11, p = 0.17), although this comparison is limited by low sample size. Nevertheless, patterns were consistent between the field and common garden, which could indicate that plants in our study are locally adapted to invest in physical defenses in response to spatial variation in herbivore abundance and damage (López-Goldar and Agrawal, 2021; Agrawal et al., 2022).

In contrast to the pattern observed in trichomes, glycoalkaloid concentrations followed a negative linear relationship across the productivity gradient where lower glycoalkaloid concentrations were produced in more productive regions. This trend occurred both in the field observations and for plants originating from across the productivity gradient in the common garden (Figs. 2, 3B, & 4B). This pattern aligns with predictions of the resource availability hypothesis, which predicts greater investment towards costly defenses under resource-limited conditions (Coley et al., 1985; Endara and Coley, 2011). Despite there being a weak correlation of glycoalkaloid production between field and common garden populations (correlation of population means: r = 0.27, p = 0.38), similar trends seen in the garden and field suggest a weak genetic basis for glycoalkaloid production. This might indicate a bottom-up, environmentally driven, selective pressure on glycoalkaloid production across plant populations rather than driven by herbivore pressure. Northern plant populations in our study system experience shorter growing seasons, and lower mean annual temperatures, conditions that could favor investment toward chemical defenses (Endara and Coley, 2011). Typically, these patterns of greater defenses in resource poor environments are found across species but have also been documented across populations especially for chemicals that are costly to produce (e.g., terpenoids; Hahn et al., 2021; Pratt & Mooney, 2013). The slight genetic basis for glycoalkaloid variation is particularly significant given that these compounds require substantial metabolic investment but provide broad-spectrum protection against herbivores through detrimental nervous system overstimulation (Wierenga and Hollingworth, 1992; Chowański et al., 2016).

Patterns of herbivore damage to plants differed between the field and common garden. In the field surveys, damage was greatest in center of the range (Fig. 3C) and was negatively correlated with glycoalkaloid concentration (Fig. 3D). In the common garden, herbivore damage was not correlated with the climate at the site of origin (Fig. 4C) but was positively correlated with SLA (Fig. 4D), which can be an indicator of leaf nitrogen and palatability. These contrasting patterns of herbivore damage in the field versus the common garden suggest that local herbivore communities likely determine different amounts of damage (Wise 2007, Wise and Rausher 2013). Although we could not robustly quantify herbivore communities at our field sites, we did observe substantial differences across the range of *S. carolinense* and in the common garden. The garden was located at the southern edge of our study gradient (Gainesville, Florida, 29.6°N), where the local herbivore community differed from the herbivore communities in the field. The three Chrysomelidae specialists, *Leptinotarsa juncta*, *Epitrix fuscula*, and *Gratiana pallidula*, often found across most of the *S. carolinense* species range, were not observed in the common garden experiment or any of the extreme edge populations (North and South; Table S1). The most common herbivore in the garden was a generalist (*Spodoptera ornithogalli*, Lepidoptera: Noctuidae). The different patterns suggest that the herbivore community is critical for understanding plant-herbivore interactions.

Additionally, although the patterns for trichomes and herbivore damage were similarly hump-shaped, there was no direct correlation between herbivory and trichomes. What this finding suggests is that trichomes may be adapting to long-term, consistently greater herbivore pressure in the center of the range (Agrawal et al., 2022; Anstett et al., 2016). Although it is possible that trichome density is responding directly to climate variation, the explanation for herbivores driving the hump-shaped relationship is more parsimonious. In contrast to trichomes, herbivory was negatively correlated with glycoalkaloid concentrations in the field, but no pattern was found in the common garden (Table 1). This might suggest that there is a defensive function of glycoalkaloids against specialist herbivores seen in the field, but not the generalist herbivores in the common garden that may select plants based on nutrition (Fig. 4D). Additionally, there was slightly more herbivory pressure in the common garden (12.1%) versus the field (9.2%), which might have contributed to a different association between glycoalkaloids and herbivory in the field versus the common garden. This shift in plant herbivore-defense relationships across different traits, where trichomes appear to be driven by herbivory whereas glycoalkaloids appear to be driven by bottom-up forces, underscores the complexity in determining defense effectiveness and evolution.

Here we demonstrate that plant defenses adapt differently across a productivity gradient, where trichome production appears to be influenced by herbivore pressure and glycoalkaloid production might be driven by climate productivity. The persistence of trichome and glycoalkaloid production trends across a productivity gradient in the field and plants derived from across a productivity gradient, indicates evolutionary adaptation to varying biotic and abiotic conditions. Overall, these findings suggest that different defensive traits within a single species are likely to follow different ecological and evolutionary trends, which could point to why plant defenses trends across a plant species range remain mixed.

## Acknowledgments

Thanks to Rebecca Molina, Emma Hair and Lucia Navia who assisted with data collection. Thanks to Monica Paniagua Montoya and Alvin Roosevelt Diamond for providing us with *Solanum carolinense* population location prospects. This study was funded by the NSF grant # DEB-1901552 to JLM and PGH and USDA McIntire-Stennis FLA-ENY-006018.

## Author contributions

J.E.H.: conceptualization; data curation; formal analysis; investigation; methodology; project administration; visualization; writing original draft, review and editing. E.S.C.: conceptualization; methodology; review and editing. V.R.C: methodology; review and editing. J.L.M.: conceptualization; funding acquisition; methodology; reviewing and editing. P.G.H.: conceptualization; data curation; formal analysis; funding acquisition; resources; investigation; methodology; supervision; review and editing.

## Data availability

Data and code are currently archived on Github: https://github.com/Plant-herbivory-interaction-lab/Herbivory-and-plant-defense-patterns-across-environmental-gradients.git, and while also be available on Dryad upon manuscript acceptance.

## Supplemental Materials

**Figure S1.**
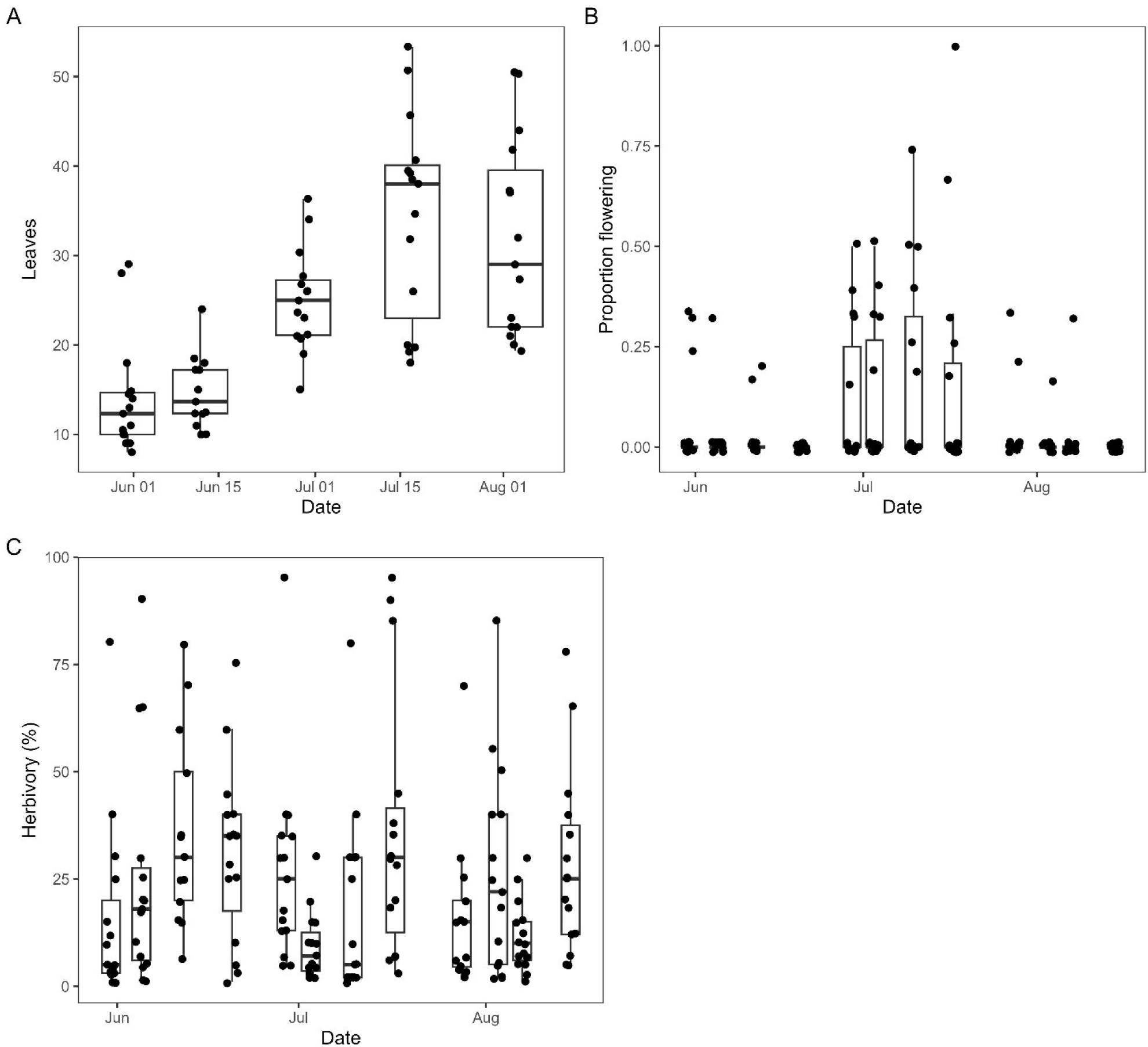
Graphs of A) leaves, B) proportion of plants flowering, and C) herbivory over time in the common garden. Data points indicate population averages at a given time point.

**Table S1.**
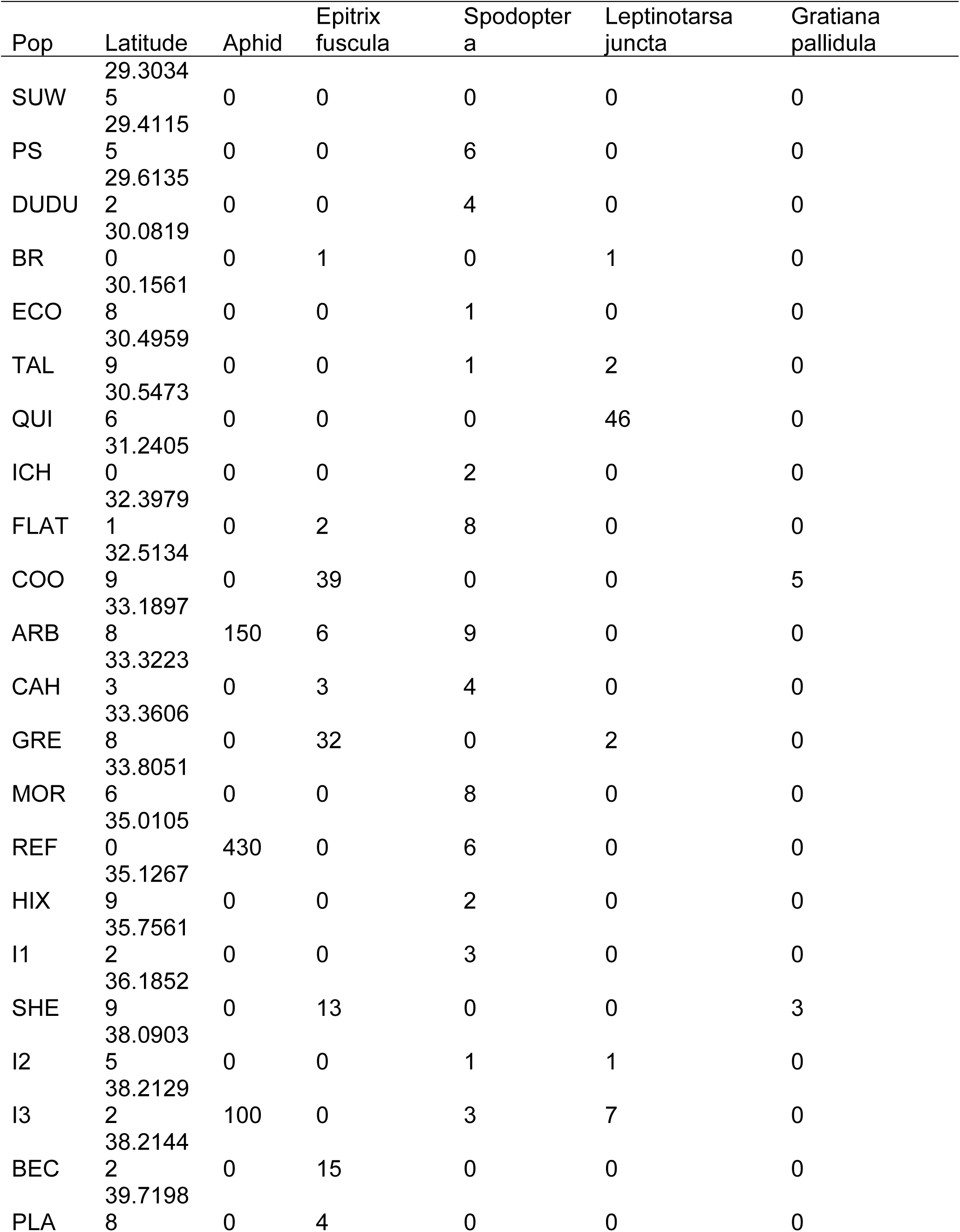
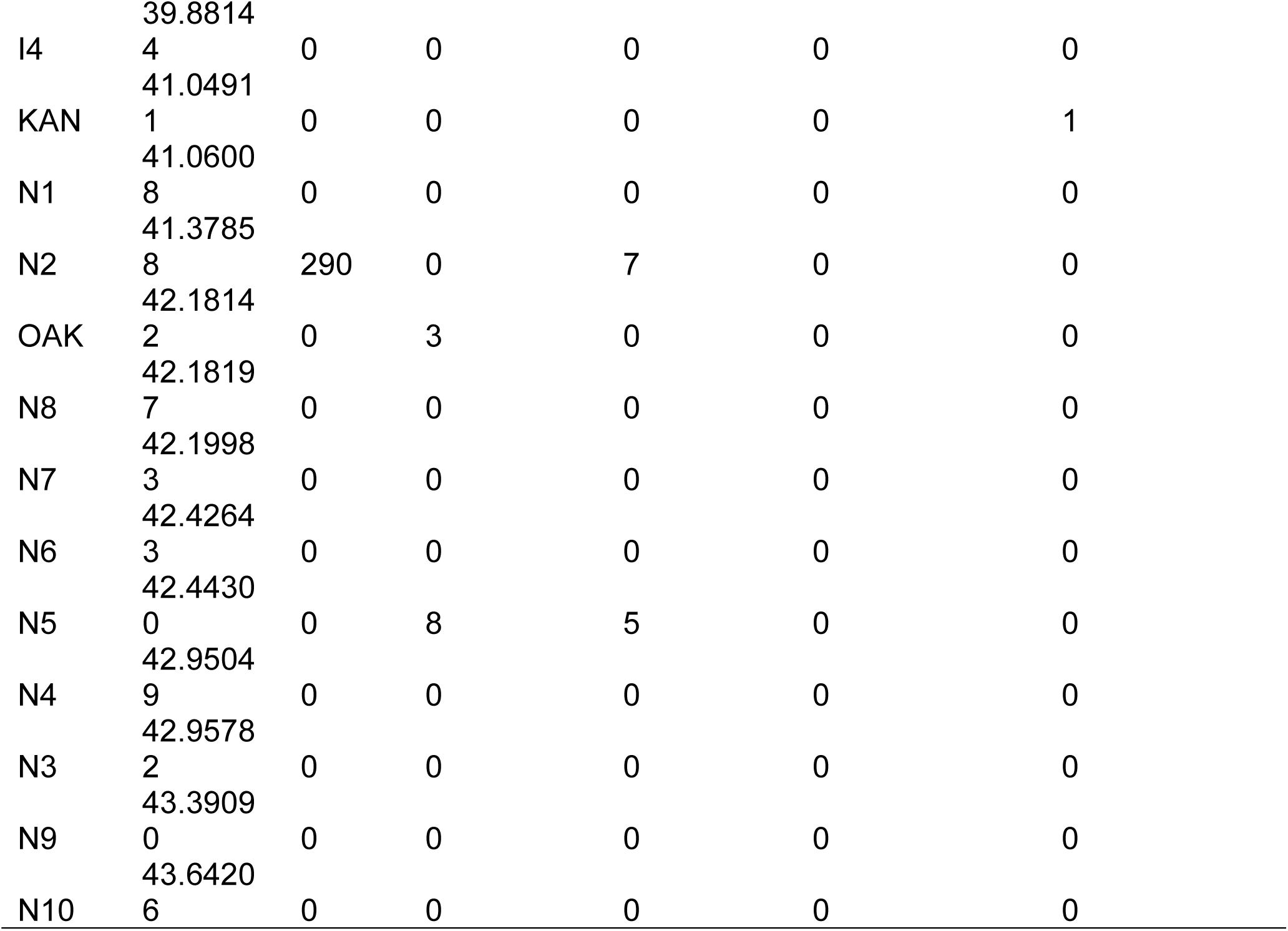
Number of insects observed during surveys of *Solanum carolinense* populations.

**Table S2.** Direct and indirect effect estimates of climate on herbivore damage derived from the field and common garden pSEM models.

| Predictor | Effect Type | Estimate |
| --- | --- | --- |
| **Field** |  |  |
| Climate PC1 | Direct | 0.305 |
| Climate PC1 | Indirect (via Conc) | -0.035 |
| Climate PC1 | Total | 0.278 |
| Climate PC1 (sq) | Direct | -0.446 |
| Climate PC1 (sq) | Indirect (via Trichomes) | 0.022 |
| Climate PC1 (sq) | Total | -0.424 |
| **Garden** |  |  |
| Climate PC1 | Direct (ns) | -0.112 |
| Climate PC1 | Indirect (via Conc, ns) | -0.005 |
| Climate PC1 | Total | -0.117 |
| Climate PC1 (sq) | Direct (ns) | 0.096 |
| Climate PC1 (sq) | Indirect (via Trichomes, ns) | -0.019 |
| Climate PC1 (sq) | Total | 0.077 |

